# Assessing the pathogenic potential of less common *Salmonella* enterica serotypes circulating in the Thai pork production chain

**DOI:** 10.1101/2022.06.27.497844

**Authors:** Thanaporn Eiamsam-ang, Pakpoom Tadee, Ben Pascoe, Prapas Patchanee

## Abstract

*Salmonella* is a frequent zoonotic foodborne pathogen, with swine and pork meats the most common source of human infection. In Chiang Mai and Lamphun Province in northern Thailand, there has been a high prevalence of salmonellosis for over a decade. Infection is usually with several dominant *S. enterica* serotypes, including serotypes Rissen and Monophasic Typhimurium. However, several less common serotypes also contribute to disease. Whole genome sequencing of 43 of these less common *S. enterica* serotypes isolated from the pork production chain through 2011-2014 were used to evaluate their genetic diversity and virulence potential. *Salmonella* contamination at local retail markets represented cross-contamination from multiple sources, including decontaminated foodstuff. Previous studies have highlighted the importance of host cell adhesion, invasion and intracellular survival for the development of clinical salmonellosis. We screened our dataset for known virulence genes and antimicrobial resistance genes, identifying at least 10 antimicrobial resistance genes in all isolates. These results indicate that these less common *S. enterica* serotypes also pose a significant public health risk. Our findings support the need for appropriate surveillance of food products going to market to reduce public exposure to highly pathogenic, multi-drug resistant *Salmonella*. Surveillance throughout the pork production chain would motivate stakeholders to reinforce sanitation standards and help reduce the risk of salmonellosis in humans.

## 1 Introduction

*Salmonella* is recognized as a prevalent bacterial-zoonotic pathogen that causes acute foodborne illness in humans and is a global public health concern (Chen et al., 2013). In the United States, approximately 1.35 million people suffer from salmonellosis with 26,500 hospitalizations and 420 deaths reported annually (CDC, 2021). In Southeast Asia, *Salmonella* has consistently contaminated the production chain for a decade, suggesting that eradication of this disease will indeed be complicated (Sinwat et al., 2016; Trongjit et al., 2017). According to the Bureau of Epidemiology of Thailand’s annual surveillance report for 2018, *Salmonella* is the most frequently detected pathogen causing food poisoning in hospitalized patients (Bureau of Epidemiology, 2018). Swine has been recognized as the one of the important *Salmonella*’s carriers, where the bacteria can multiply in the digestive tract and can be spread to other steps of the production chain via feces (Rostagno & Callaway, 2012). Furthermore, pork has been reported to be an important source of *Salmonella* contamination especially in the retail market which is the predisposing factor of salmonellosis in human (Patchanee et al., 2016).

Investigations have been undertaken to quantify the prevalence of *Salmonella* throughout the pork production chain in Chiang Mai, Thailand. These studies have reported a high prevalence of *Salmonella* isolated from swine farms as 30.56% (Tadee et al., 2014), and up to 42% in retail pork circulating in the Chiang Mai municipality area (Patchanee et al., 2016). These finding suggest that the burden of *Salmonella* has been continuing and substantial for a decade. All aspects of the production chain have been contaminated, from the farm-slaughterhouse-retail market, which is likely related to the levels of sanitation and hygienic at each step of production (Savall et al., 2016; Chen et al., 2019). *Salmonella* Rissen and Monophasic *Salmonella* Typhimurium have been the most common serotypes isolated in Chiang Mai and Lamphun provinces for more than a decade and their epidemiology investigated using molecular typing methods (Prasertsee et al., 2019; Patchanee et al., 2020). In contrast, many of the less common serotypes, such as *S. enterica* serotypes Anatum, Panama, Stanley and Give have not been studied further.

Whole genome sequencing-based methods are now being used to identify transmission networks and assess the genetic relatedness of the clonal or closely related strains during an outbreak (Oakeson et al., 2017). Core genome MLST provides high-resolution data and reveals the precise relatedness within the species by comparing allelic variation to equivalent loci in other isolates (Pearce et al., 2020). The emergence of antimicrobial resistance (AMR) in foodborne pathogens particularly *Salmonella* species has posed a significant threat to public health (Ferri et al., 2017). Indiscriminate antimicrobial use in the livestock industry has been identified as a driver for multidrug-resistant (MDR) organisms, which can be spread to humans through the food chain (Van et al., 2015; Nhung et al., 2016). In addition, the pathogenic potential of *Salmonella* has been linked to expression of virulence genes. Adhesion and invasion genes are essential virulence genes, which when expressed allow colonization of infecting *Salmonella* of the host cell (Wang et al., 2020). Expression of genes required for growth and replication within the host are then required for adequate nutrient uptake (Saha et al., 2019). Furthermore, the virulence genes that are encoded for resistance to host defense and resistance to antimicrobial peptide are the one of the important mechanisms to allowing the chronic infection of *Salmonella* (Kintz et al., 2015; Jajere S. M., 2019). Whole genome sequence-based techniques can help characterize isolates according to their putative pathogenic potential, by identifying known antimicrobial resistance genes (AMRs) and virulence-related genes (Campioni & Faicao, 2012).

In this study, we sequenced isolates from these less common *Salmonella* serotypes isolated from farms, slaughterhouses, and retail markets in Chiang Mai and Lamphun provinces between 2011 and 2014. We characterize the genetic diversity of these isolates and compare their genetic relatedness throughout the production process to assess feasibility of transmission. In addition, we characterize the antimicrobial resistance genes and virulence genes of each isolate in order to better understand the potential risk of these less common lineages. These data will help assess public health risk and inform public health guidance and prevention strategies for minimize *Salmonella* contamination in the study area.

## 2 Materials & Methods

### 2.1 Bacterial isolates

A total of 43 *S. enterica* isolates from the pork production chain in Chiang Mai and Lamphun municipality area were included in this study. These samples were collected from three steps of the pork production chain, including farms (n=17), slaughterhouses (n=16) and retail markets (n=10). Classification by serotype including *Salmonella enterica* serotypes Stanley (n=12), Typhimurium (n=10), Panama (n=6), Give (n=6), Krefeld (n=2), Kedougou (n=2), Anatum (n=1), Agona (n=1), Lexington (n=1), Newport (n=1) and Yoruba (n=1). Isolates were serotyped according to the WHO National *Salmonella* and *Shigella* Center Laboratory (NSSC) in Nonthaburi, Thailand using the slide agglutination method and serotypes were assigned according to the Kauffmann-White scheme (Brenner et al., 2000). Typing details for each isolate is shown in **Table 1**.

**Table 1:** Origin and characteristic of *S. enterica* isolates tested recovered from farms, slaughterhouses and retail markets during the period of 2011-2014.

| ID | Serotype | Year | Source | Province |
| --- | --- | --- | --- | --- |
| 8440 | S. Typhimurium | 2011 | Farm | Chiang Mai |
| 8448 | S. Typhimurium | 2011 | Farm | Chiang Mai |
| 8453 | S. Typhimurium | 2011 | Farm | Lamphun |
| 8454 | S. Typhimurium | 2011 | Farm | Lamphun |
| 8455 | S. Typhimurium | 2011 | Farm | Chiang Mai |
| 8456 | S. Typhimurium | 2011 | Farm | Lamphun |
| 8457 | S. Typhimurium | 2011 | Farm | Lamphun |
| 8458 | S. Typhimurium | 2011 | Farm | Lamphun |
| 8464 | S. Typhimurium | 2011 | Farm | Lamphun |
| 8512 | S. Panama | 2011 | Farm | Lamphun |
| 8513 | S. Panama | 2012 | Farm | Lamphun |
| 8516 | S. Give | 2012 | Farm | Chiang Mai |
| 8517 | S. Give | 2012 | Farm | Chiang Mai |
| 8518 | S. Give | 2012 | Farm | Chiang Mai |
| 8522 | S. Stanley | 2012 | Farm | Lamphun |
| 8527 | S. Stanley | 2011 | Farm | Lamphun |
| 8529 | S. Stanley | 2012 | Farm | Lamphun |
| 8459 | S. Typhimurium | 2013 | Slaughterhouse | Lamphun |
| 8508 | S. Panama | 2013 | Slaughterhouse | Chiang Mai |
| 8509 | S. Panama | 2013 | Slaughterhouse | Chiang Mai |
| 8510 | S. Panama | 2013 | Slaughterhouse | Chiang Mai |
| 8511 | S. Panama | 2013 | Slaughterhouse | Chiang Mai |
| 8514 | S. Give | 2013 | Slaughterhouse | Lamphun |
| 8515 | S. Give | 2013 | Slaughterhouse | Chiang Mai |
| 8519 | S. Stanley | 2013 | Slaughterhouse | Lamphun |
| 8520 | S. Stanley | 2013 | Slaughterhouse | Lamphun |
| 8521 | S. Stanley | 2013 | Slaughterhouse | Lamphun |
| 8524 | S. Stanley | 2013 | Slaughterhouse | Lamphun |
| 8525 | S. Stanley | 2013 | Slaughterhouse | Lamphun |

| **ID** | **Serotype** | **Year** | **Source** | **Province** |
| --- | --- | --- | --- | --- |
| **8526** | S. Stanley | 2013 | Slaughterhouse | Chiang Mai |
| **8528** | S. Stanley | 2013 | Slaughterhouse | Chiang Mai |
| **8530** | S. Stanley | 2013 | Slaughterhouse | Chiang Mai |
| **8531** | S. Stanley | 2013 | Slaughterhouse | Chiang Mai |
| **8425** | S. Anatum | 2014 | Market | Chiang Mai |
| **8431** | S. Krefeld | 2014 | Market | Chiang Mai |
| **8434** | S. Kedougou | 2014 | Market | Chiang Mai |
| **8436** | S. Krefeld | 2014 | Market | Chiang Mai |
| **8438** | S. Newport | 2014 | Market | Chiang Mai |
| **8872** | S. Lexington | 2014 | Market | Chiang Mai |
| **8877** | S. Kedougou | 2014 | Market | Chiang Mai |
| **8878** | S. Agona | 2014 | Market | Chiang Mai |
| **8879** | S. Yoruba | 2014 | Market | Chiang Mai |
| **8883** | S. Give | 2014 | Market | Chiang Mai |

### 2.2 Whole genome sequencing

The DNA of all 43 *S. enterica* isolates were extracted using QIAamp DNA mini kits (Qiagen, Crawley, UK). The Nextera XT DNA Library Preparation Kit was used for the library preparation according to the manufacturer’s instructions (Illumina, Cambridge, UK). *Salmonella* genomes were sequenced as short reads using an Illumina MiSeq 300 bp paired-end sequencer (Illumina, Cambridge, UK). Short reads were filtered, trimmed with TRIMMOMATIC (Bolger et al., 2014), and assembled *de novo* with SPAdes software (version 3.8.0, using the-careful command) (Bankevich et al., 2012). The average number of contigs was 341 (range: 89-2365) for an average total assembled sequence size of 4,918,241 bp (range: 4,169,306-5,234,359). The average N50 was 38,380 (range: 2249-128,256) and the average GC content was 52.2% (range: 52.0-52.6). Short read data are available on the NCBI Sequence Read Archive, associated with BioProjects PRJNA573746 and PRJNA419926 (https://www.ncbi.nlm.nih.gov/sra).

### 2.3 Core genome multilocus sequence typing (cgMLST)

Whole genome sequencing data of all 43 *S. enterica* isolates were uploaded to a Bacterial Isolate Genome Sequence Database (BIGSdb) and the genome comparator tool (constructed for the and provided by pubMLST) (https://pubmlst.org/bigsdb?db=pubmlst_salmonella_isolates&page=plugin&name=GenomeComparator) used to assess gene presence among the isolates. The genome comparator tool analyzes all selected isolates using the EnteroBase *Salmonella* database’s cgMLST scheme, which considers a total of 3002 loci. (Pearce et al., 2020)

To expand our collection, we included an additional 61 *S. enterica* isolates from public repositories: Enterobase *Salmonella enterica* WGS database (https://enterobase.warwick.ac.uk/species/index/senterica) (Warwick Medical School**; Supplementary Table S1**). These additional genomes included isolates from different source reservoirs in Thailand during the period of 2001 through 2016. Isolates were typed to *Salmonella enterica* serotypes Stanley (n=8), Typhimurium (n=21), Panama (n=2), Give (n=1), Anatum (n=8), Kedougou (n=9), Agona (n=7) and Lexington (n=5); from animals (2 wild animals, and 1 poultry), food (38 pork, 6 frozen seafood, 5 spices, 3 foods and 2 vegetables), and human (n = 4). A minimum spanning tree of all isolates was constructed, based on advanced cluster analysis for categorical data of allelic number for cgMLST using Bionumerics version 7.6 (Applied Maths, Ghent, Belgium).

### 2.4 Virulence genes and antimicrobial resistance genes investigation

We used the RASTtk algorithm (Brettin et al., 2015) to annotate whole genome sequencing data of all isolates via the PATRIC v3.6.12 annotation server using default parameters. (Wattam et al., 2017) (https://patricbrc.org/app/Annotation). The Virulence Factor Database (VFDB; database version 2019) (Liu et al., 2019) was used to define the presence of known virulence genes including adhesion effector, invasion effectors, intracellular survival effectors and toxin-producing genes. Antimicrobial resistance gene presence was explored using the Comprehensive Antibiotic Resistance Database (CARD; database version 2020) (Alcock et al., 2020). The antimicrobial resistance genes (AMR genes) related to the expression of aminoglycoside, beta-lactam, trimethoprim, fluoroquinolone, fosfomycin, macrolide, macrolide-lincosamide-streptogramin B, peptide, phenicol, sulfonamide, and tetracycline were investigated. The dendrogram of the virulence genes and antimicrobial resistance genes investigation were constructed using the unweighted pair group method with arithmetic mean (UPGMA) algorithms according to the cluster analysis of categorical values of genes concluded in this study.

## 3 Results

### 3.1 *S. enterica* serotypes differ in their ability survive through the pork production chain

From the minimum spanning tree analysis, all *S. enterica* isolates tested were divided into 4 major clusters with 9 additional singleton isolates (**Figure 1A**). According to the source of origin, isolates from farms and slaughterhouses were grouped together into the similar clusters. Conversely, most isolates recovered from retail markets (9/10) did not cluster with farm or slaughterhouse isolates. There was a single cluster that was comprised of isolates from all three sources of *S. enterica* isolates, supporting persistence of this serotype (*S. enterica* serotype Give) from the swine farm to slaughter and contamination of retail pork products (**Figure 1B**). This cluster was comprised solely of isolates collected in Chiang Mai and Lamphun province during the period of 2012-2014.

**Figure 1:**
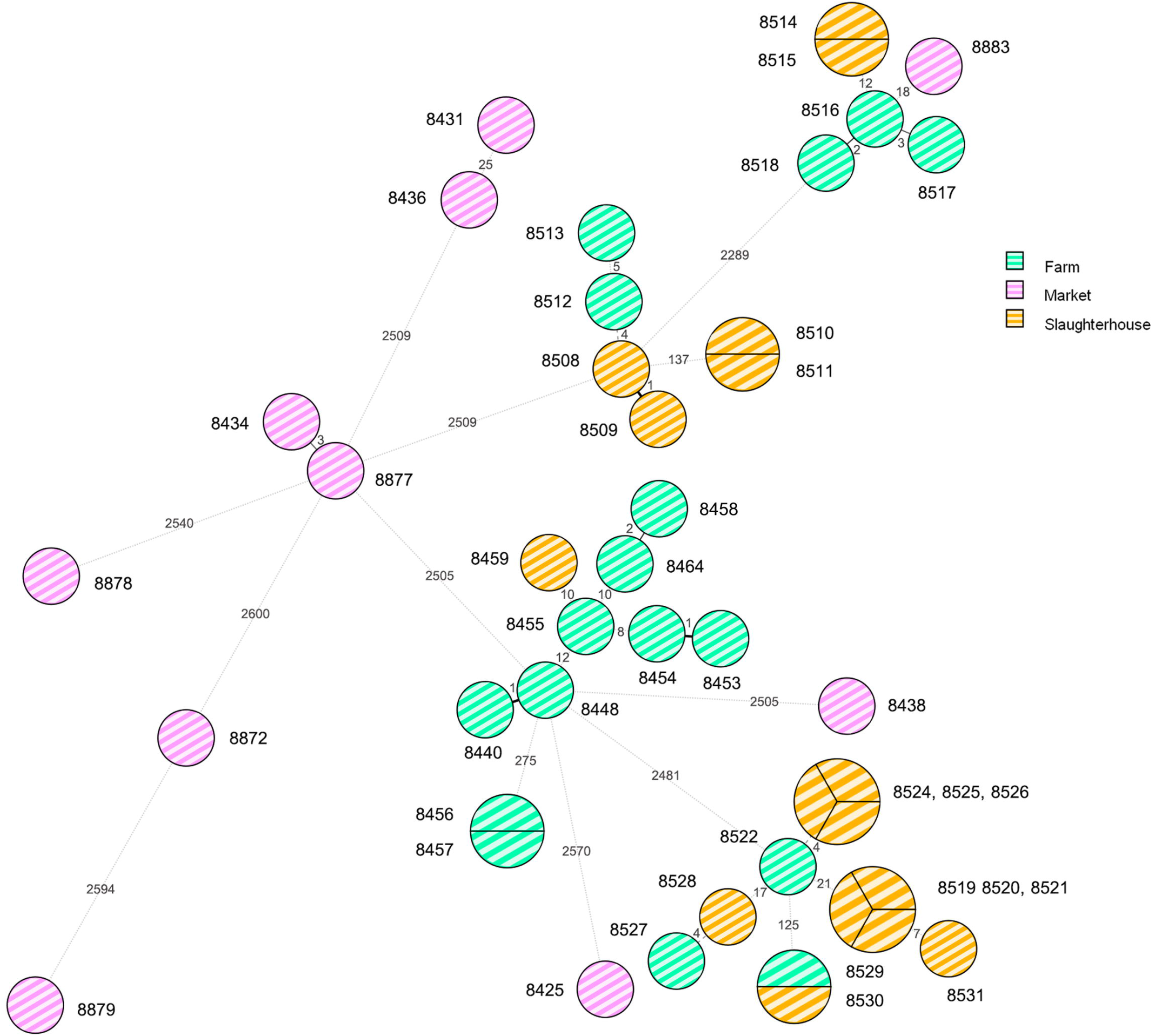
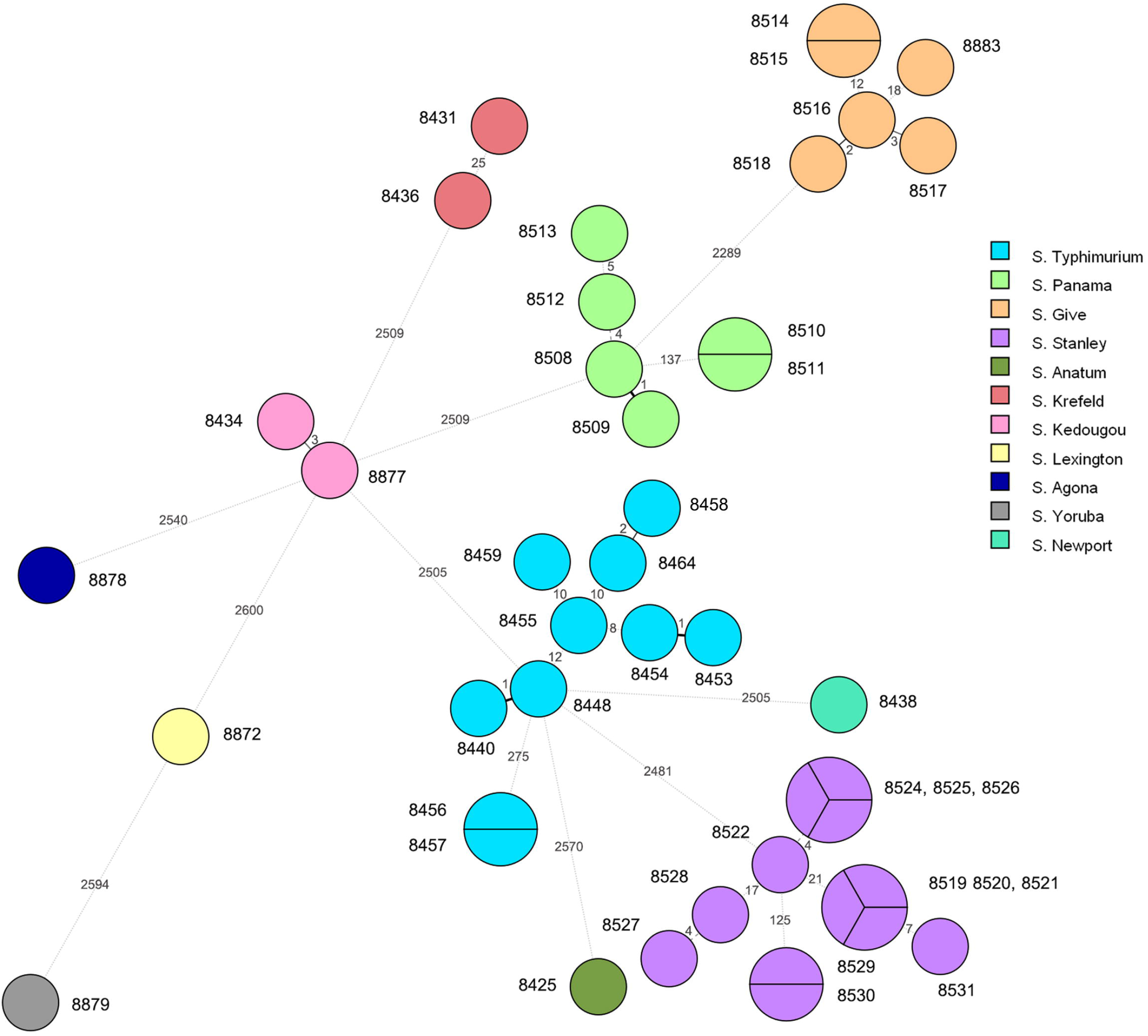
The minimum spanning tree (MST) analysis of *Salmonella* isolates recovered from the pork production chain. Each isolate was grouped according to the loci different of cgMLST scheme. (A): Node color coding: green color, yellow color and pink color represent the *Salmonella* isolates recovered from farm, slaughterhouse and retail market, respectively. (B): Node color coding: each color represents each serotype of *Salmonella* isolates.

Some *Salmonella* isolates shared the same cgMLST profiles, including serotype Typhimurium isolates ID 8456 and 8457 – both recovered from farms in Lamphun province in the period of 2011; six serotype Stanley isolates (IDs: 8519, 8520, 8521, 8524, 8525, & 8526) recovered from slaughterhouses in Chiang Mai and Lamphun during the period of 2013; and two serotype Panama isolates (IDs: 8510 & 8511) from Chiang Mai which were recovered from slaughterhouse in 2013 (**Figure 1**). In addition, some clonal isolates were able to persist between different production steps and time periods, including two serotype Stanley isolates (IDs: 8529 and 8530): collected from a farm in Lamphun in 2012, and a slaughterhouse in Chiang Mai in 2013, respectively. Two others clonal serotype Give isolates were collected from slaughterhouses in different provincial areas during 2013 (IDs: 8514 & 8515; **Figure 1**).

Notwithstanding *S. enterica* serotype Rissen and the monophasic Typhimurium, which are the most common serotypes identified in the northern Thai pork production chain, *S. enterica* serotypes Typhimurium, Stanley and Panama were the predominant serotypes recovered from farms and slaughterhouses. These *Salmonella* isolates were found in Chiang Mai and Lamphun Province during the period of 2011 through 2013. Other serotypes, including *S. enterica* serotypes Agona, Anatum, Krefeld, Kedougou, Lexington, Newport, and Yoruba were not grouped into any clusters and were isolated from sources in the retail markets in the Chiang Mai municipality area. *S. enterica* serotype Give was the only serotype that was found in all 3 different steps during 2012-2014\ (**Figure 1B**).

### 3.2 Not all high-risk *S. enterica* isolates collected in the retail markets were from pork

To better understand the genetic relatedness of isolates in our collection, we constructed a minimum spanning tree based on cgMLST profiles of our collected isolates (n=43), compared with all publicly available genomes from Thailand (n=61; **Supplementary Table S1**). In total, 104 *S. enterica* isolates were clustered according to their cgMLST profiles by their source of origin **(Figure 2)**, and there was a distinction in isolates from the farms and slaughterhouses and those collected in the retail markets. Isolates collected from the retail markets demonstrated greater diversity in cgMLST profiles (6 cgMLST clusters) and serotypes (7 Serotypes). Four cgMLST clusters and 4 serotypes overlapped between the farms and slaughterhouses **(Supplementary Figure S1)**. Many of the isolates collected in Chiang Mai municipality markets were of unique cgMLST profiles and serotypes that were not present in any of our pork production samples, instead grouping with clusters from other Thai food products, including spices, food, frozen seafood and pork. Although pork products are a frequent source of salmonellosis infection, they are not the only high-risk food product available in the retail markets. The limited number of clinical *S. enterica* isolates that were publicly available (n=4) clustered (same/similar cgMLST profile) predominantly with isolates identified in the pre-harvest steps of pork production (n=3), with only a single isolate clustering with isolates identified at retail markets **(Figure 2)**. Given the low number of isolates compared it’s difficult to draw robust conclusions but does suggest that isolates from the pork production industry are able to persist and pose a public health risk.

**Figure 2:**
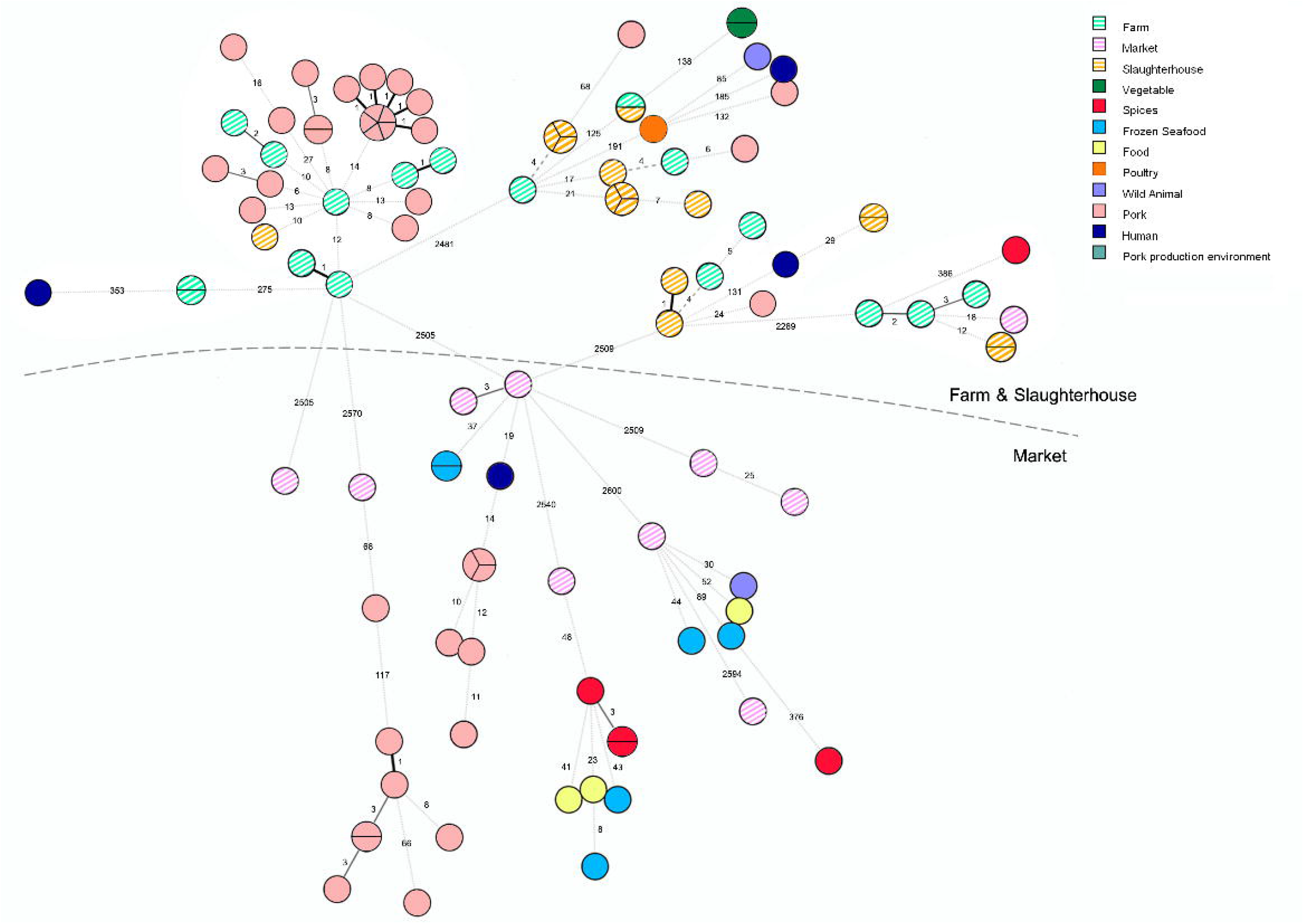
The minimum spanning tree (MST) analysis of 43 *Salmonella* isolates recovered from pork production chain (striped nodes) and additional 61 *Salmonella* isolates circulating in Thailand. Node color coding were representing the sources of the *Salmonella* isolates.

### 3.3 Virulence gene profiling of *S. enterica* from the pork production chain

Known virulence genes were identified in all isolates through nucleotide comparisons with the VFDB. All isolates carried the virulence genes encoding for host cell adhesion (*csg, fim, mis, sin*), host cell invasion (*che, flg, fli, inv, mot, omp, org, prg, sic, sif, sip, slr, sop, spa, spt, ssa, ssc, sse*) and intracellular survival (*ent, fep, gmh, iro, mgt, kds, mig*). The virulence genes were harbored in the *Salmonella* isolates regardless of serotype, production steps, year, and geographical area. However, serotype Give isolates seemed to contain alternative host adhesion genes (*fae* and *shd*) in place of the more common *lpf* and *rat* genes. We also identified *cdt* genes in these serotypes Give isolates (and serotype Panama), which are typically associated with *S. typhi* and encode for toxin production. This is particularly alarming as serotype Give isolates were able to persist through the pork production chain, and therefore these highly virulent genes were identified in isolates from farms, slaughterhouses, and retail markets (**Figure 3**).

**Figure 3:**
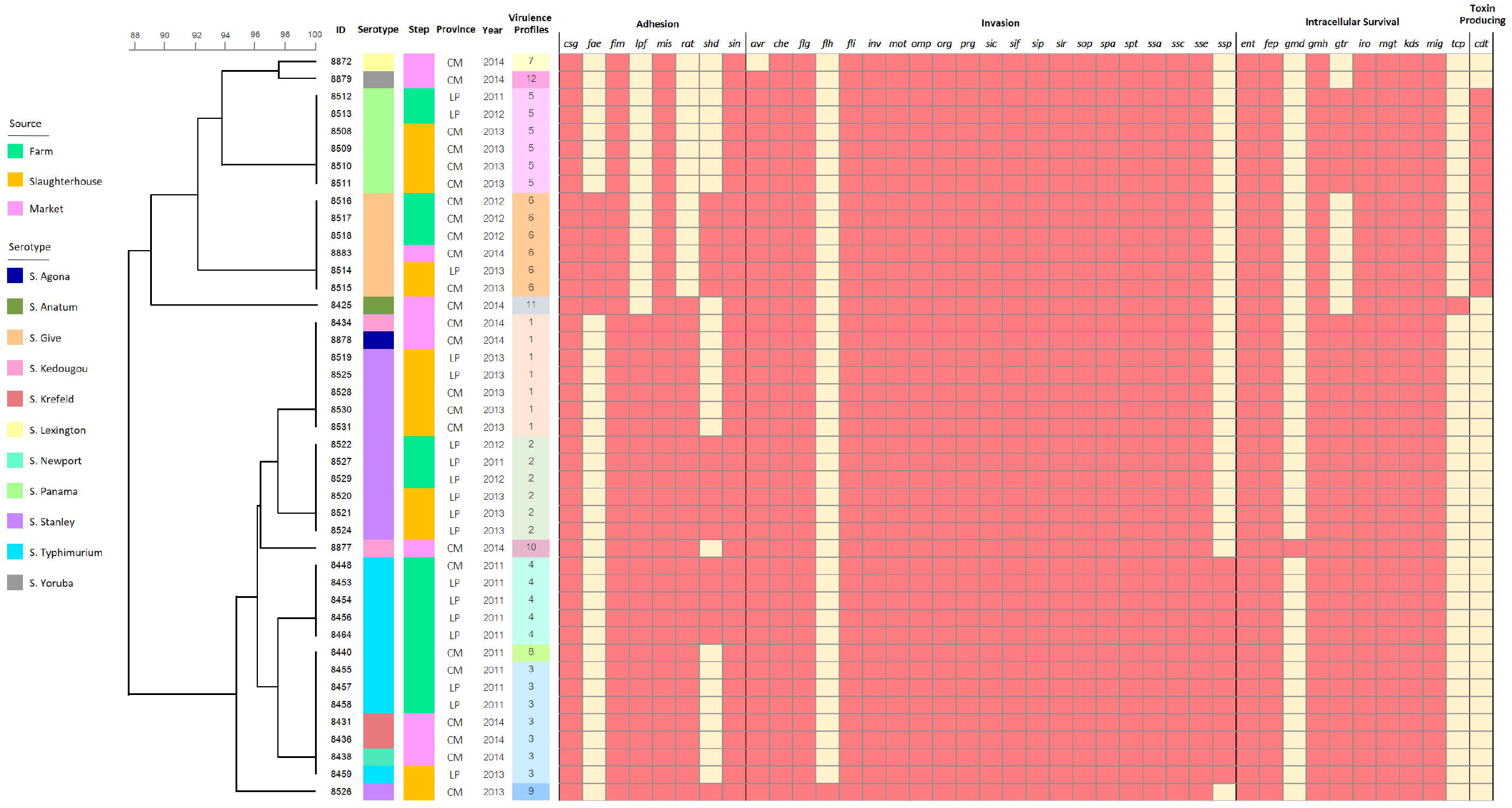
Binary heatmap analysis of virulence genes harbored in the *Salmonella* isolates recovered from 3 production steps circulating in Chiang Mai and Lamphun municipality area during the period 2011-2014.

Isolates were characterized according to presence of each of the 40 identified virulence genes and classified into 12 virulence profiles. Profiles 1 and 3 (16.27%) were the most frequently encoded profile, followed by profiles 2, 5 and 6 with 13.95% of frequency. Several genes were found only in specific sources, resulting in unique virulence profiles, including virulence profiles 9 (with *flh* genes isolated from the slaughterhouse), profiles 10 and 11 (with *gmd* and *tcp* genes isolated from the retail markets, respectively). There was also diversity in virulence profiles among serotypes and the different steps of the production chain. In another words, there are the less concordance between serotyping results and virulence profiles (**Figure 3**).

An intersection analysis of virulence genes demonstrated that 37 virulence genes were shared among the three steps of pork production. Many *Salmonella* isolates collected from the retail markets had the most virulence genes (39/40 virulence genes), followed by *Salmonella* isolates recovered from slaughterhouses and farms (38 and 37 virulence genes were identified, respectively) (**Supplementary Figure S2**).

### 3.4 Widespread multidrug resistance in isolates from all steps of the pork production chain

According to the heatmap analysis, all the *Salmonella* isolates in this study carried at least one antimicrobial resistance gene (ARG), and all of them were multi-drug resistant *Salmonella* – demonstrating putative resistance to three or more antimicrobial classes. Similar to the distribution of virulence genes, ARGs were predominantly lineage dependent - with isolates from the same serotype sharing similar ARG content, even across different isolation sources, time and geographical area. In total, ten antimicrobial resistance genes were harbored in all *Salmonella* isolates, including *acrD, gyrA, mfd, parC, parE, glpT, murA, bacA, pmrC*, and *pmrE*, which contribute towards aminoglycoside, fluoroquinolone, fosfomycin, and peptide resistance, while CTX-M-14 and *erm* (42), which encoded for beta-lactam and macrolide-lincosamide-streptogramin B resistance, was not found in this study (**Figure 4**).

**Figure 4:**
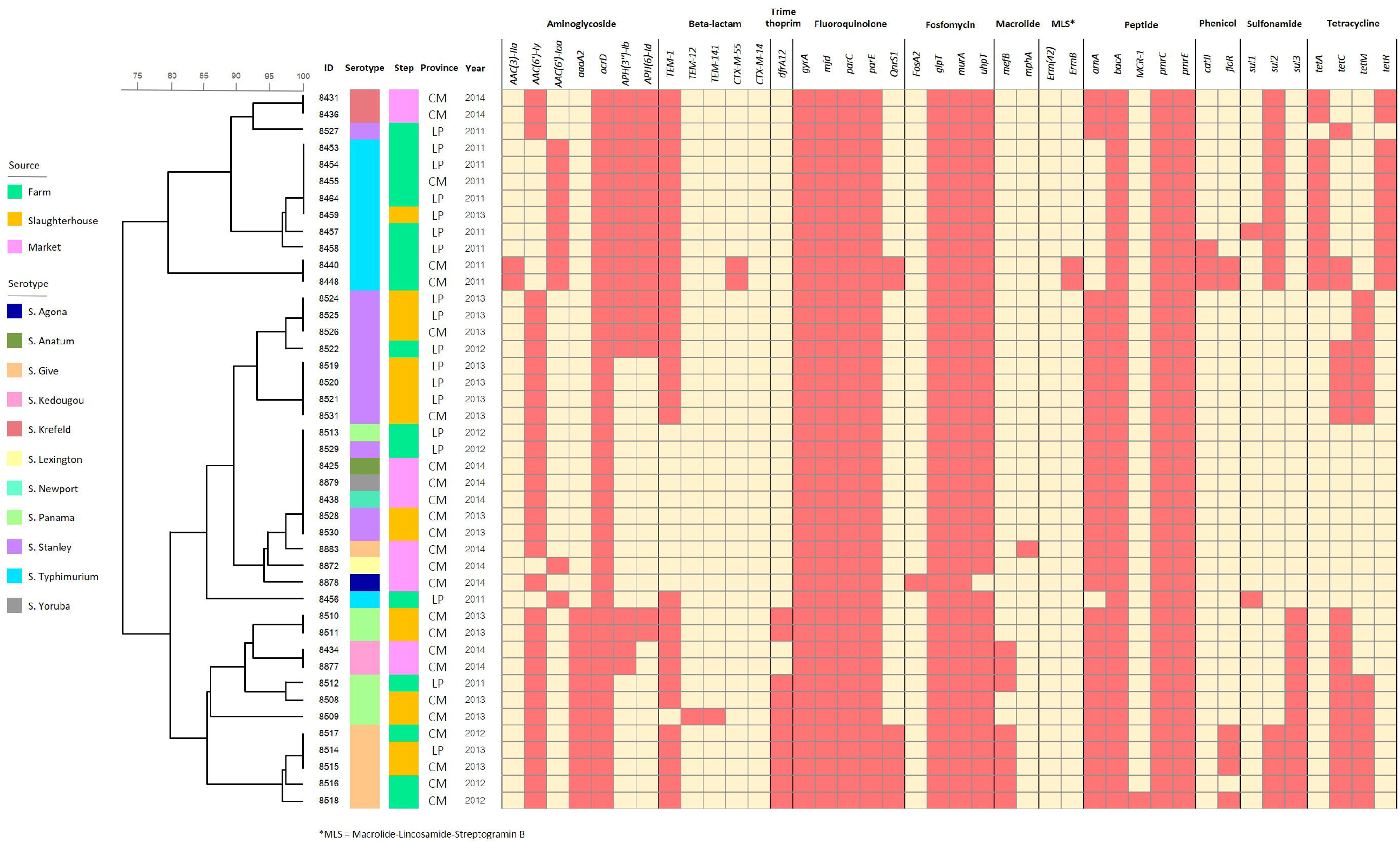
Binary heatmap analysis of antimicrobial resistance genes harbored in the *Salmonella* isolates recovered from 3 production steps circulating in Chiang Mai and Lamphun municipality area during the period 2011-2014.

According to intersection analysis, twenty-four antimicrobial resistance genes were shared throughout all three pork production steps. Additionally, four antimicrobial resistance genes (*drfA12, QnrS1, florR* and *tetM*) were identified in isolates sampled from farms and slaughterhouses, which are linked to trimethoprim, fluoroquinolone, phenicol and tetracycline, respectively. The *Salmonella* isolates recovered from farms harbored the highest number of antimicrobial resistance genes (34/40 ARGs), followed by the *Salmonella* isolates recovered from slaughterhouses and retail markets (30 and 26 ARGs identified, respectively). Some ARGs were identified in specific sources of *Salmonella* isolates, including *AAC (3)-IIa, CTX-M-55, ErmB, MCR-1, catII* and *sul1* genes in farm isolates, while *TEM-12* and *TEM-141* genes were only found in slaughterhouses. Furthermore, *FosA2* and *mphA* were only found in *Salmonella* isolates that were on the retail markets. (**Supplementary Figure S3**)

## 4 Discussion

The most common *Salmonella* serotypes implicated in widespread contamination of the pork production chain in northern Thailand are monophasic Typhimurium and serotype Rissen. Previous genomics studies have focused on these common serotypes, with less common serotypes overlooked (Prasertsri et al., 2019; Patchanee et al., 2020). In this study we focus on these less common serotypes to investigate their genetic diversity and virulence potential. Isolates originating from farm and slaughterhouse were closely related, with isolates from serotypes Typhimurium, Panama, and Stanley collected from both production steps. This is consistent with previous studies suggesting that these serotypes are common at the pre-harvest level. (Sanguankiat et al., 2010; Niyomdecha et al., 2016). Together, these findings support the feasibility of *Salmonella* spreading from the farms to slaughterhouses via live animals (Rostagno & Callaway, 2012; Savall et al., 2016).

*S. enterica* serotype Give was the only serotype found at all three steps of the pork production chain. This serotype has previously been found in Thailand from diverse sources of food-animal production, including poultry, pork and other food products, such as chili powder (Wang et al., 2015; Phongaran et al., 2019). Another study implicated this serotype as the causative agent of salmonellosis with a splenic abscess in a male patient who had travelled to southern Thailand and consumed raw minced pork (Girardin et al., 2006). Despite this, many isolates found primarily on the farms and in the slaughterhouses apparently pose no direct impact on consumers, i.e., they are not able to survive (or out compete other strains) to contaminate pork products sold at market. However, there is evidence that *S. enterica* serotypes found predominantly at the pre-harvest level can cause infection in workers at the operational level (Sringam et al., 2017). Estimates suggest that as many as 43% of workers are colonized by *Salmonella*, compared to an overall prevalence of *Salmonella* contamination in farmed pigs of 52%. This suggests that transmission between pigs and humans is very common in the farm environment, with *S. enterica* serotype Typhimurium implicated as the primary infecting agent (Punpanich et al.,2012). *S. enterica* serotypes Stanley and Panama were also among the most common serotypes isolated from farms and slaughterhouses and have been recovered from the stool samples of salmonellosis patients in other studies (Pulford et al., 2019). The *S. enterica* isolates that we collected from retail markets were more diverse and represented serval *Salmonella* serotypes.

Of the 10 samples we collected from the retail markets, we identified 8 different serotypes. There was little overlap between serotypes found in retail market isolates and those sampled on the farms or slaughterhouses. It is likely that this diversity, in both phenotypic and genotypic characteristics are due to the wider sources of contamination of retail products than the pork production industry (Chen et al., 2019). We identified a single isolate from the retail market samples that was from *S. enterica* serotype Anatum, which is typically one of the most common serotypes found in pork products in Thailand (Padungtod & Kaneene, 2006) - sometimes considered the 3^rd^ most common serotype isolated from pork in Asia (Ferrari et al., 2019). Other serotypes, including Agona, Kedougou and Lexington are grouped in clusters with isolates from frozen seafood and spices in Thailand. Contamination with *Salmonella* from uncooked seafood products sold at market has been estimated at the rate of 21% (Woodring et al., 2012). However, these serotypes have been identified from multiple agricultural products and retail food products, including seafood, meat, spices, and herbs (Zhao et al., 2006). Spices such as clove, oregano, black pepper, red chili, and pepper powder have become important sources of *Salmonella* contamination, and have been implicated in foodborne outbreaks in several countries, e.g., contaminated fresh basil in Denmark; and herbal tea in Germany (Koch et al., 2005; Pakalniskiene et al., 2009). Cross-contamination between foodstuffs at the retail point cannot be discounted due to the remaining of *Salmonella* on the food contact surface, which difficult to eradicate from the retail environment and facilities. (De Cesare et al., 2003; Campos et al., 2019).

Although some virulence genes were detected at low frequency, many of the virulence genes we identified were found in nearly all isolates, including those associated with host cell adhesion, invasion and intracellular survival (Jajere et al., 2019). Alarmingly, we also were able to identify cytolethal distending toxin genes (*cdt*) in a small number of isolates from serotypes Give and Panama. Typically, *cdt* genes are found in highly virulent *Salmonella* Typhi isolates, although there is precedence for their presence in non-typhoidal *Salmonella* (Haghjoo & Galan, 2004). Previous work has identified *cdt* genes in non-typhoidal *Salmonella* serotypes Javiana, Montevideo, and Oranienburg, and were implicated in their higher capability to persist and cause infection (Miller & Wiedmann, 2016). Virulence genes were found in all isolates at all steps of the pork production chain, timescale, or geographical area. Isolates from the same serotype shared similar virulence profiles, which transcended sampling source. This suggests that all isolates that we collected had the potential to colonize and infect humans, whether through direct consumption by consumers of retail food products or exposure of workers on the farms and slaughterhouses (Poonchareon et al., 2019).

The global rise in AMR pathogens is a significant public health risk and forecasts on death tolls resulting from AMR infections are shocking – with an estimated death toll of 10 million people by 2050, if no action is taken (Balouiri et al; 2016). Fluoroquinolone resistant *Salmonella* are among the WHO’s high priority organisms for development of new antibiotics (WHO, 2017). In our collection, all 43 isolates were multidrug resistant and resistant to at least three antimicrobial classes. Previous studies have identified widespread dissemination of MDR *Salmonella* throughout the pork production chain, with high prevalence (98%) in Thailand (Sinwat et al., 2016). Multi-drug resistant *Salmonella* have also been observed in the neighboring countries, including Laos (98.4%) and Cambodia (52%) (Trongjit et al., 2017). Isolates carried at least different 10 ARGs, associated with aminoglycoside, fluoroquinolone, fosfomycin, and peptide resistance. All antimicrobials where putative resistance was identified are widely used in veterinary practice, especially in swine production – where farmed swine are thought to consume more antimicrobials than any other livestock animal (Van et al., 2015; Nhung et al., 2016). The highest number of ARG was found in farm isolates, followed by the slaughterhouses and then the retail markets. Clearly, there is a strong selective pressure imposed by incongruous antimicrobial usage, evidenced here and in other studies by a high prevalence of AMR organisms (not just *Salmonella*) in industrially farmed swine (Harada & Asai, 2010; Tadee et al., 2015). Our findings support the bleak WHO outlook where a rise in fluoroquinolone resistant organisms continues to erode the efficacy of antimicrobials for clinical cases and there is an urgent need to either curtail this rise; or develop novel antimicrobials (Lee et al., 2009).

The lack of strong regulation and indiscriminate use of antimicrobials has promoted dissemination of ARGs in the Thai food chain (Lekagul et al., 2021). As a result, the Department of Livestock Development, Ministry of Agriculture and cooperatives have implemented a ban on the use of any antimicrobials as a growth promoter in animal feed to combat antimicrobial resistance in livestock animals. Even though the government agency has already issued some policies for reducing the antimicrobial resistance pathogens’ occurrence in the livestock section, some antimicrobial agents, including aminoglycoside and fluoroquinolone, are still available for treatment at the farm level (Nuangmek et al., 2021). Our findings indicate that multidrug resistant *Salmonella* are still a problem in the pork production chain, and larger scale surveillance studies are required.

## 5 Conclusion

In the Chiang Mai and Lamphun Municipality areas, *Salmonella* circulating in pork production is hazardous to humans in every step along the production chain. All of these isolates contained the necessary virulence genes for the pathogenicity of salmonellosis. In addition, the *Salmonella* isolates in this study were the multi-drug resistance *Salmonella* which harmful to the public health worldwide. In terms of epidemiological knowledge, there is less information about the less common serotypes of *Salmonella* in this study area. This study reveals the possible common ancestors of each *Salmonella* serotype and can provide additional information about the evidence of *Salmonella*’s cross-contamination at the pre-harvest level. Furthermore, the whole genome sequence-based analysis can substantiate the possibility of *Salmonella* contamination from other agricultural products to the pork at the retail level. Additional information from this study would motivate all stakeholders to be aware of and pay attention to the reinforcement of standardized sanitation throughout the pork production chain in order to eradicate and reduce the risk of *Salmonella* contamination, which affects public health worldwide.

## 6 Author Contributions

TE conceived and designed the experiments, performed the experiment, analyzed the data, prepared figures and/or tables, and approved the final draft. PT conceived and designed the experiments, prepared figures and/or tables, authored or reviewed drafts of the paper, and approved the final draft. BP conceived and designed the experiments, performed the experiment, analyzed the data, and approved the final draft. PP conceived and designed the experiments, authored or reviewed drafts of the paper, and approved the final draft.

## 8 Acknowledgements

All high-performance computing was performed on MRC CLIMB (supported by MRC grants MR/L015080/1 and MR/T030062/1). Finally, the authors would like to express our appreciation to our colleagues at Chiang Mai University for noteworthy contributions and partial funding support.

## 10 Data Availability Statement

The datasets analyzed for this study can be found in the NCBI associated with the Bio project: PRJNA573746 and PRJNA419926.

Supplementary Figure S1: The minimum spanning tree (MST) analysis of 43 *Salmonella* isolates recovered from pork production chain (striped nodes) and additional 61 *Salmonella* isolates circulating in Thailand. Node color coding were representing the serotypes of the *Salmonella* isolates.

Supplementary Figure S2: The Venn diagram of intersection analysis of virulence genes among different steps of pork production chain. The Venn diagram represent the number of unique and shared virulence genes in 43 *Salmonella* isolates recovered from farms, slaughterhouses and retail markets.

Supplementary Figure S3: The Venn diagram of intersection analysis of antimicrobial resistance genes among different steps of the pork production chain. The Venn diagram represents the number of unique and shared antimicrobial resistance genes in 43 *Salmonella* isolates recovered from farms, slaughterhouses, and retail markets.

Supplementary Table S1: Origin and characteristic of *S. enterica* isolates originating from various sources in Thailand during the period of 2001 through 2016 acquired from public repositories (Enterobase *Salmonella* database).

