## Supplementary material for "Assessing the pathogenic potential of less common *Salmonella* enterica serotypes circulating in the Thai pork production chain": 968695_Table_1.DOCX

Supplementary Table S1: Origin and characteristic of *S. enterica* isolates originating from various sources in Thailand during the period of 2001 through 2016 acquired from public repositories (Enterobase *Salmonella* database)

| Accession Number | Serotype | Year | Source |
| --- | --- | --- | --- |
| SRS845256 | S. Stanley | 2001 | Vegetable |
| SRS1357486 | S. Stanley | 2001 | Wild Animal |
| SRS1297595 | S. Lexington | 2002 | Wild Animal |
| SRS528748 | S. Panama | 2003 | Human |
| SRS528775 | S. Typhimurium | 2003 | Human |
| SRS845347 | S. Stanley | 2003 | Vegetable |
| SRS735136 | S. Lexington | 2004 | Frozen Seafood |
| SRS798489 | S. Agona | 2005 | Frozen Seafood |
| SRS932622 | S. Lexington | 2005 | Spices |
| SRS978996 | S. Lexington | 2008 | Frozen Seafood |
| SRS528717 | S. Kedougou | 2008 | Human |
| SRS528767 | S. Stanley | 2008 | Human |
| SRS850553 | S. Agona | 2008 | Spices |
| SRS1417505 | S. Lexington | 2009 | Food |
| SRS856111 | S. Agona | 2009 | Spices |
| SRS856077 | S. Agona | 2009 | Spices |
| SRS1011312 | S. Kedougou | 2010 | Frozen Seafood |
| SRS1011304 | S. Kedougou | 2010 | Frozen Seafood |
| SRS528080 | S. Agona | 2011 | Frozen Seafood |
| SRS580801 | S. Agona | 2012 | Food |
| SRS1380650 | S. Anatum | 2013 | Pork |
| SRS1416354 | S. Anatum | 2013 | Pork |
| SRS1380653 | S. Kedougou | 2013 | Pork |
| SRS1380782 | S. Kedougou | 2013 | Pork |
| SRS1380685 | S. Panama | 2013 | Pork |
| Accession Number | **Serotype** | **Year** | **Source** |
| SRS1380649 | S. Stanley | 2013 | Pork |
| SRS1416362 | S. Stanley | 2013 | Pork |
| SRS1380659 | S. Typhimurium | 2013 | Pork |
| SRS1380686 | S. Typhimurium | 2013 | Pork |
| SRS1416350 | S. Typhimurium | 2013 | Pork |
| SRS1416357 | S. Typhimurium | 2013 | Pork |
| SRS1416358 | S. Typhimurium | 2013 | Pork |
| SRS1416359 | S. Typhimurium | 2013 | Pork |
| SRS1416360 | S. Typhimurium | 2013 | Pork |
| SRS1416361 | S. Typhimurium | 2013 | Pork |
| SRS1416363 | S. Typhimurium | 2013 | Pork |
| SRS1416364 | S. Typhimurium | 2013 | Pork |
| SRS1416367 | S. Typhimurium | 2013 | Pork |
| SRS1416368 | S. Typhimurium | 2013 | Pork |
| SRS1416386 | S. Typhimurium | 2013 | Pork |
| SRS1416387 | S. Typhimurium | 2013 | Pork |
| SRS1416388 | S. Typhimurium | 2013 | Pork |
| SRS1416389 | S. Typhimurium | 2013 | Pork |
| SRS1416413 | S. Typhimurium | 2013 | Pork |
| SRS1416373 | S. Anatum | 2014 | Pork |
| SRS1416374 | S. Anatum | 2014 | Pork |
| SRS1416375 | S. Anatum | 2014 | Pork |
| SRS1416378 | S. Anatum | 2014 | Pork |
| SRS1416380 | S. Anatum | 2014 | Pork |
| SRS1416383 | S. Anatum | 2014 | Pork |
| SRS1416377 | S. Kedougou | 2014 | Pork |
| SRS1416379 | S. Kedougou | 2014 | Pork |
| SRS1416381 | S. Kedougou | 2014 | Pork |
| SRS1416382 | S. Kedougou | 2014 | Pork |
| SRS1416376 | S. Stanley | 2014 | Pork |
| SRS1353138 | S. Typhimurium | 2014 | Pork |
| Accession Number | **Serotype** | **Year** | **Source** |
| SRS1353148 | S. Typhimurium | 2014 | Pork |
| SRS1416384 | S. Typhimurium | 2014 | Pork |
| SRS621678 | S. Give | 2014 | Spices |
| SRS6951023 | S. Agona | 2016 | Food |
| SRS4318372 | S. Stanley | 2016 | Poultry |
