## Supplementary figures and images for "Assessing the pathogenic potential of less common *Salmonella* enterica serotypes circulating in the Thai pork production chain"

### 968695_Image_1.JPEG

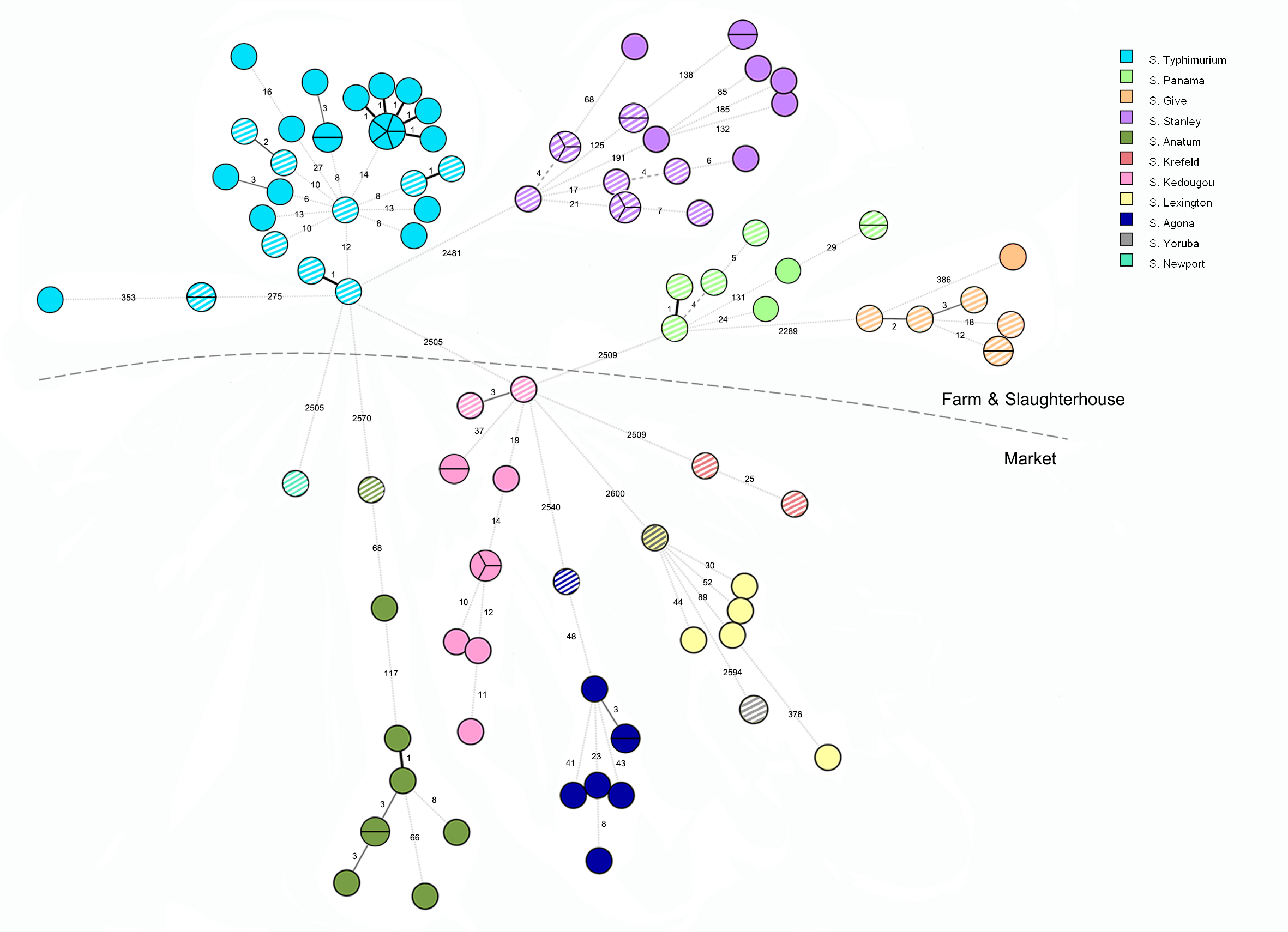

### 968695_Image_2.JPEG

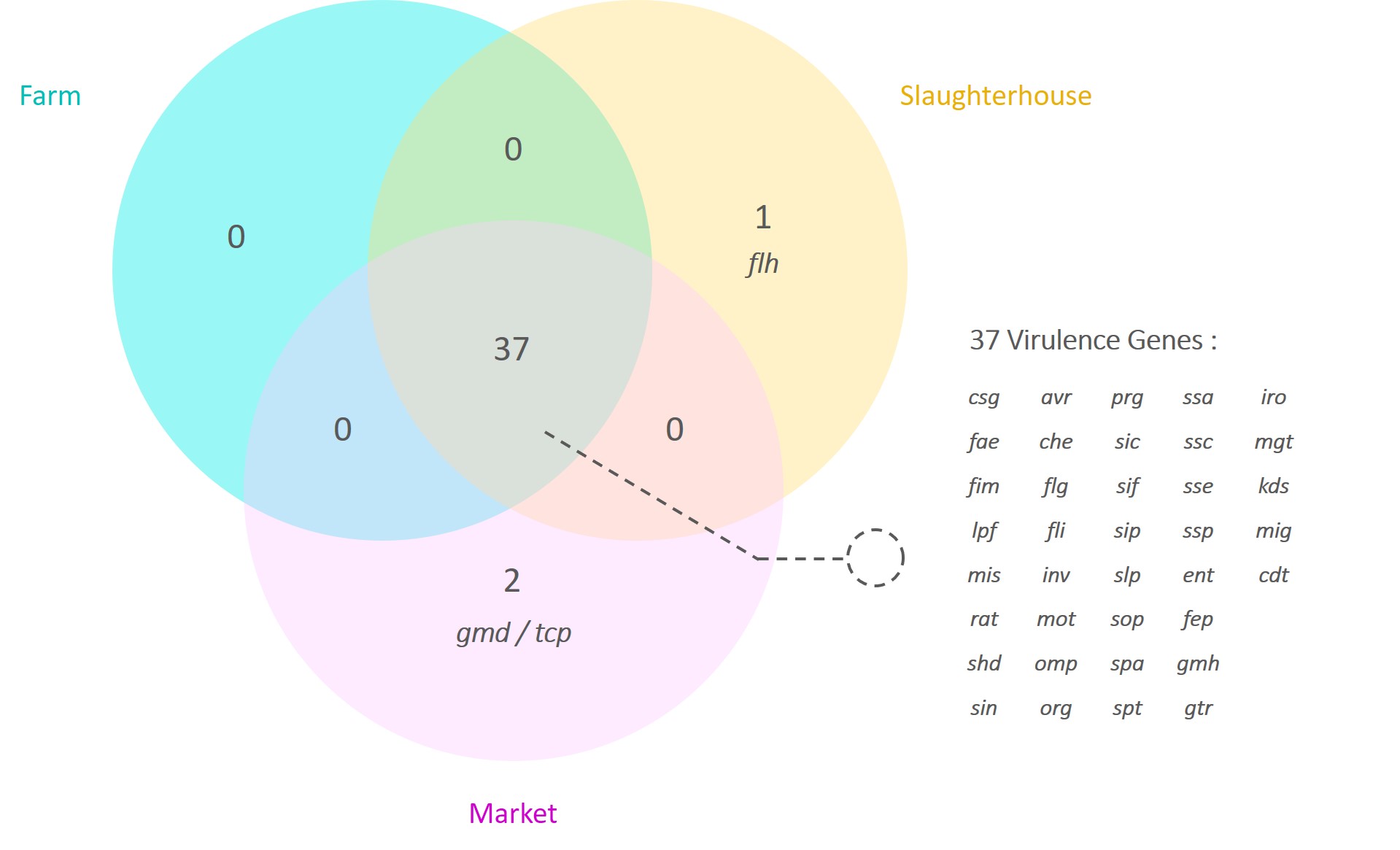

### 968695_Image_3.JPEG

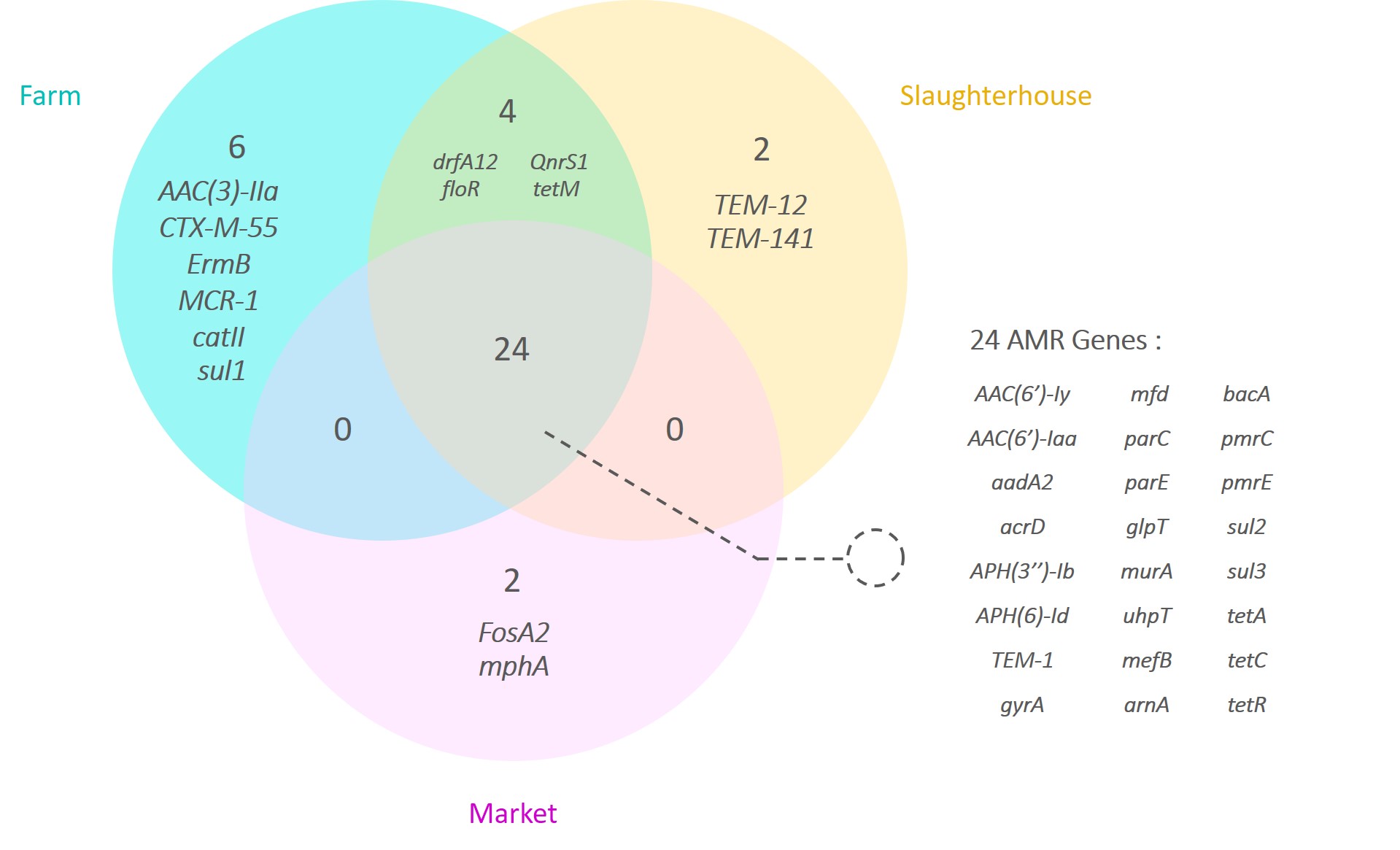
